# In-cell structures of a conserved supramolecular array at the mitochondria-cytoskeleton interface in mammalian sperm

**DOI:** 10.1101/2021.02.16.431372

**Authors:** Miguel Ricardo Leung, Riccardo Zenezini Chiozzi, Marc C. Roelofs, Johannes F. Hevler, Ravi Teja Ravi, Paula Maitan, Min Zhang, Heiko Henning, Elizabeth G. Bromfield, Stuart C. Howes, Bart M. Gadella, Albert J.R. Heck, Tzviya Zeev-Ben-Mordehai

## Abstract

Mitochondria-cytoskeleton interactions modulate cellular physiology by regulating mitochondrial transport, positioning, and immobilization. However, there is very little structural information defining mitochondria-cytoskeleton interfaces in any cell type. Here, we use cryo-focused ion beam milling-enabled cryo-electron tomography to image mammalian sperm, where mitochondria wrap around the ciliary cytoskeleton. We find that mitochondria are tethered to their neighbors through inter-mitochondrial linkers and are anchored to the cytoskeleton through ordered arrays on the outer mitochondrial membrane. We use subtomogram averaging to resolve in-cell structures of these arrays from three mammalian species, revealing they are conserved across species despite variations in mitochondrial dimensions and cristae organization. We find that the arrays consist of boat-shaped particles anchored on a network of membrane pores whose arrangement and dimensions are consistent with voltage dependent anion channels. Proteomics and in-cell cross-linking mass spectrometry suggest that the conserved arrays are composed of glycerol kinase-like proteins. Ordered supramolecular assemblies may serve to stabilize similar contact sites in other cell types where mitochondria need to be immobilized in specific subcellular environments, such as in muscles and neurons.

## Introduction

In many cell types, mitochondria collectively form a dynamic network whose members divide, fuse, and communicate with one another (Glancy et al., 2015; Viana et al., 2020; Vincent et al., 2017). Through interactions with the cytoskeleton, mitochondria are transported – sometimes across large distances – and positioned in response to dynamic stimuli (Fenton et al., 2021; Moore and Holzbaur, 2018). Interactions with the cytoskeleton can also restrain mitochondria to specific subcellular locations. In neurons, axonal mitochondria can be immobilized by interactions with the microtubule or actin cytoskeletons (Chen and Sheng, 2013; Gutnick et al., 2019; Kang et al., 2008). In cardiac and skeletal muscle, mitochondrial distribution is regulated by inter-actions with myofibrils and intermediate filaments (Milner et al., 2000; Stone et al., 2007). However, despite the prevalence of inter-mitochondria and mitochondria-cytoskeleton interactions and their integral roles in cellular function, there is very little information on the molecular architectures of these interaction sites in any cell type.

One of the most striking mitochondrial configurations occurs in amniote sperm, where mitochondria are arranged in a spiral around the axoneme, defining a region called the midpiece (Fawcett, 1970, 1975). Mitochondria are among the few organelles retained in sperm throughout their maturation, during which they otherwise lose most of their cytoplasm and organelles *en route* to becoming highly streamlined cells specialized for finding and fusing with the egg. The extensive mitochondrial sheath in amniote sperm may be an adaptation needed to power the large, long flagellum in these lineages. Variations in midpiece morphometry affect sperm motility and competitiveness (Firman and Simmons, 2010; Fisher et al., 2016), and different species rely on energy from mitochondrial respiration to different extents (Marin et al., 2003; Tourmente et al., 2015), warranting comparative studies of mitochondrial structure across species.

The core of the midpiece is the ciliary cytoskeleton, composed of the microtubule-based axoneme and accessory elements called outer dense fibers (ODFs). A poorly-characterized network of cytoskeletal filaments called the submitochondrial reticulum lies between the ODFs and the mitochondria. The submitochondrial reticulum co-purifies with the outer mitochondrial membrane (OMM), suggesting that they are intimately associated (Olson and Winfrey, 1986, 1990). Mitochondria wrap around the cytoskeleton and are in turn ensheathed by the plasma membrane. As a consequence of this arrangement, each mitochondrion has three distinct surfaces (Olson and Winfrey, 1992) – one facing the axoneme, one facing the plasma membrane, and one facing neighboring mitochondria. Thin-section electron microscopy (EM) (Olson and Winfrey, 1992) and freeze-fracture EM (Friend and Heuser, 1981) suggest that each surface is characterized by a unique membrane protein profile. Notwithstanding the insight gained from these methods, such techniques require harsh sample preparation steps that can distort fine cellular structure and limit achievable resolution (Al-Amoudi et al., 2004). As such, the molecular landscape of inter-mitochondrial and mitochondrial-cytoskeleton contacts in the sperm midpiece remains largely unexplored.

Assembly of the mitochondrial sheath occurs late in spermiogenesis and involves an intricately choreographed series of events (Ho and Wey, 2007; Otani et al., 1988). Initially, spherical mitochondria are broadly distributed in the cytoplasm. Mitochondria are then recruited to the flagellum, where they form ordered rows along the flagellar axis. Finally, mitochondria elongate and twist around the axoneme. While our understanding of the molecular details of these processes is cursory at best, studies on gene-disrupted mice have implicated a number of proteins in mitochondrial sheath morphogenesis. For instance, mice expressing mutant forms of kinesin light chain 3 (KLC3) have malformed midpieces, hinting at a role for microtubule-based transport (Zhang et al., 2012). Another example are the voltage dependent anion channels (VDACs), which are highly abundant proteins that mediate transport of metabolites, ions, and nucleotides like ATP across the OMM (Colombini, 2012). Male mice lacking VDAC3 are infertile and their sperm cells have disorganized mitochondrial sheaths (Sampson et al., 2001), so VDACs may also have unappreciated roles in mitochondrial trafficking; indeed, KLC3 binds mitochondria through VDAC2 (Zhang et al., 2012). Similarly, disrupting sperm-specific isoforms of glycerol kinase leads to gaps in the mitochondrial sheath despite proper initial alignment of spherical mitochondria (Chen et al., 2017b; Shimada et al., 2019). Mice lacking spermatogenesis-associated protein 19 (SPATA19) (Mi et al., 2015) or glutathione peroxidase 4 (GPX4) (Imai et al., 2009; Schneider et al., 2009) also have structurally abnormal mitochondria.

Here, we use cryo-focused ion beam (cryo-FIB) milling-enabled cryo-electron tomography (cryo-ET) to image the mitochondrial sheath in mature sperm from three mammalian species. We take advantage of the uniquely multi-scale capabilities of cryo-ET to unveil new aspects both of the overall organization of the mitochondrial sheath and of the molecular structures important for its assembly. We find that mitochondria are tethered to their neighbors through inter-mitochondrial linkers and to the underlying cytoskeleton through conserved protein arrays on the OMM. Subtomogram averaging revealed that these arrays are anchored on a lattice of OMM pores whose arrangement and dimensions are consistent with VDACs. Proteomics and in-cell cross-linking mass spectrometry suggest that the arrays consist of glycerol kinase (GK)-like proteins. Our data thus show that although mitochondrial dimensions and cristae architecture vary across species, the architecture of the mitochondria-cytoskeleton interface is conserved at the molecular level.

## Results

### Mitochondrial dimensions and cristae organization vary across species

We imaged the mitochondrial sheath in mature sperm from three mammalian species, namely the pig (*Sus scrofa*), the horse (*Equus caballus*), and the mouse (*Mus musculus*) **(Fig. 1)**. These species differ in terms of sperm size, motility patterns, and metabolism. To visualize the overall organization of the mitochondrial sheath, we imaged whole sperm with a Volta phase plate (VPP) (Danev et al., 2014; Fukuda et al., 2015). Neural-network based segmentation (Chen et al., 2017a) of the mitochondrial membrane allowed us to visualize mitochondrial organization in three dimensions **(Fig. 1a-f)**.

**Fig. 1.**
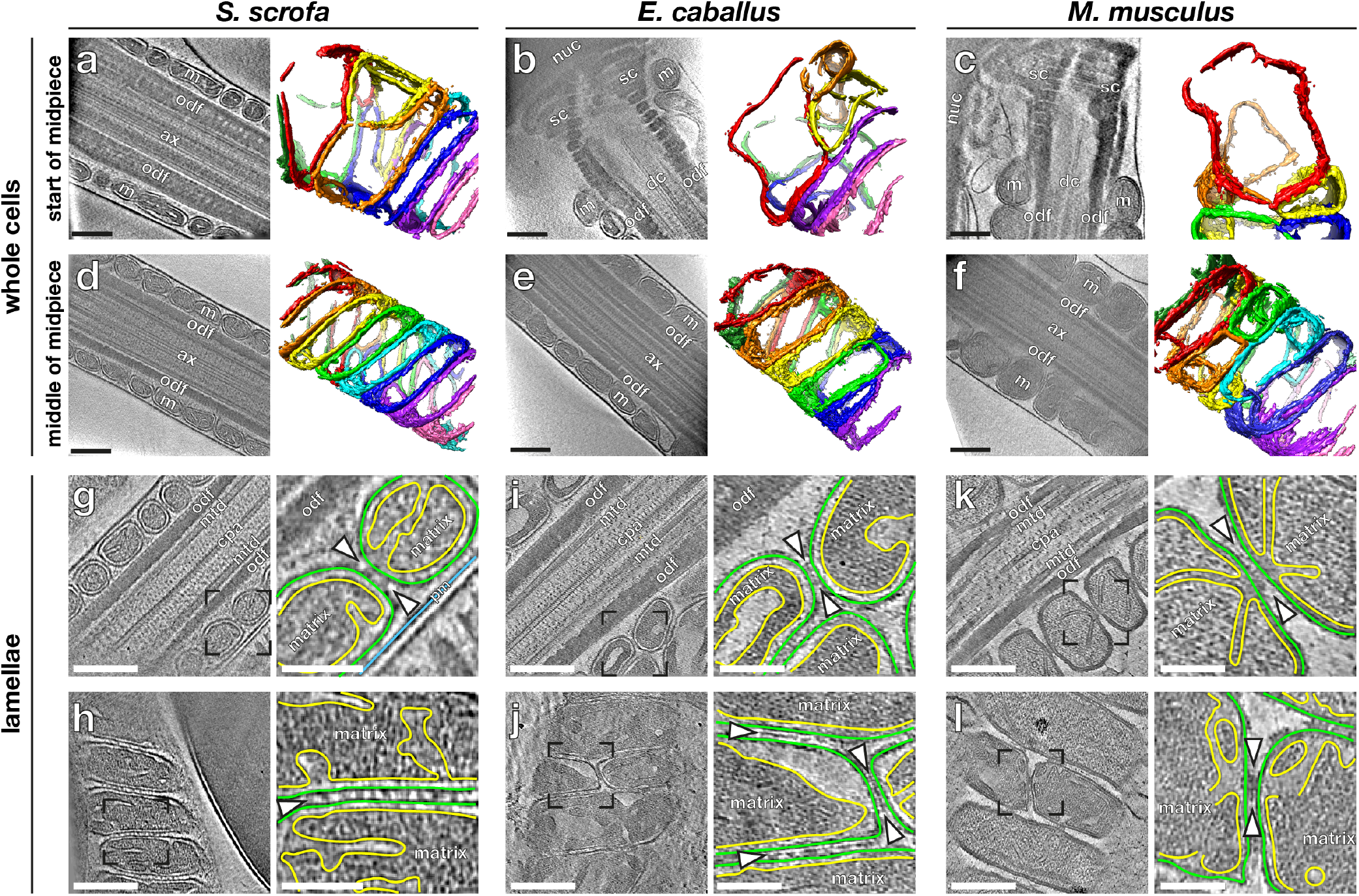
Mitochondrial dimensions and cristae organization vary across species. **(a-f)** Slices through Volta phase plate cryo-tomograms (left) and corresponding three-dimensional segmentations (right) of mitochondria from the start (a-c) or middle (d-f) of the midpiece from pig (a,d), horse (b,e), and mouse (c,f) sperm. **(g-l)** Slices through cryo-tomograms of FIB-milled pig (g,h), horse (i,j), and mouse (k,l) sperm midpieces. Right panels show digital zooms of the regions boxed out in the left panels. The outer mitochondrial membrane is traced in green, the inner mitochondrial membrane in yellow, and the plasma membrane in blue. Arrowheads indicate inter-mitochondrial linker complexes. **Labels:** nuc – nucleus, sc – segmented columns, m – mitochondria, odf – outer dense fibers, dc – distal centriole, ax – axoneme, mtd – microtubule doublets, cpa – central pair apparatus, pm – plasma membrane. **Scale bars:** (a-l) left panels – 250 nm, (g-l) right panels – 100 nm.

To investigate variations in mitochondrial width along the midpiece, we first measured the width of each mitochondrion at multiple points along its length. We then divided mitochondria into groups based on their positions along the midpiece, as measured by their distance from the head **(Fig. S1)**. The midpiece is ~10 μm long in both pig and horse sperm, but ~20 μm long in mouse sperm, so each group represents ~2 μm in the pig and the horse and ~4 μm in the mouse. We found that mouse sperm mitochondria are ~1.5 times wider than pig and horse sperm mitochondria overall **(Fig. S1a)**. In all three species studied, most mitochondria in the middle (~60%) of the midpiece are crescent-shaped tubes **(Fig. 1d-f)** with consistent widths along their lengths **(Fig. S1b)**. Mitochondria at the proximal end of the midpiece are larger than their more distal counterparts **(Fig. 1a-c, S1a)**. Moreover, proximal mitochondria have more variable shapes, evidenced by greater variation in their widths at different point along their lengths **(Fig. S1b)**. Because mitochondria wrap around the axoneme, variations in mitochondrial dimensions both across species and along the proximodistal axis of the flagellum affect the overall diameter and rigidity of the midpiece, likely fine-tuning the hydrodynamics of sperm motility.

To visualize the internal organization of sperm mitochondria in a near-native state, we imaged sperm thinned by cryoFIB milling **(Fig. 1g-l)**. This revealed unexpected diversity in the internal ultrastructure of mitochondria across mammalian species, especially in terms of cristae morphology. Horse sperm mitochondria have an expanded intermembrane space and a condensed matrix **(Fig. 1i-j)**. Mouse sperm mitochondria have an expanded matrix, with a narrow intermembrane space and thin cristae **(Fig. 1k-l)**. Pig sperm mitochondrial morphology is intermediate **(Fig. 1g-h)**, and although the mitochondrial matrix was dense, we could identify individual complexes that resembled ATP synthase on cristae of FIB-milled mitochondria **(Fig. S2a-b)**, which was confirmed by subtomogram averaging **(Fig. S2b’)**.

Inter-species differences in cristae morphology correlate with measurements of matrix volume relative to mitochondrial volume **(Fig. S2d)**. In this regard, horse sperm mitochondria resemble “condensed” mitochondria, which correlate with higher rates of oxidative activity in a number of different cell types, including developing germ cells, neurons, and liver (Hackenbrock, 1968; De Martino et al., 1979; Perkins and Ellisman, 2011). Indeed, horse sperm are dependent on oxidative phosphorylation (Davila et al., 2016), whereas pig (Marin et al., 2003) and mouse sperm (Mukai and Okuno, 2004; Odet et al., 2013) are thought to rely largely on a glycolytic mechanisms.

### Inter-mitochondrial junctions are associated with linker complexes

Mitochondria are closely packed within the mitochondrial sheath, but it is unclear whether or how individual organelles communicate with their neighbors. To address this, we imaged inter-mitochondrial junctions captured in FIB-milled sperm lamellae. We observed transmitochondrial cristae alignment in mouse sperm **(Fig. 1k-l)**, but not in pig or in horse sperm **(Fig. 1g-j)**. Trans-mitochondrial cristae alignment has also been observed in muscle tissue of various organisms, and is proposed to mediate electrochemical coupling between adjacent mitochondria (Picard et al., 2015). To our knowledge, this is the first time this phenomenon has been observed in mature sperm from any lineage. It is particularly curious, however, that trans-mitochondrial cristae alignment in sperm is species-specific.

We found that inter-mitochondrial junctions are characterized by novel inter-mitochondrial linker complexes in all three species **(arrowheads in Fig. 1g-l, Fig. S2c)**. These inter-mitochondrial linkers span the 8-nm distance between the outer membranes of neighboring mitochondria. In mouse sperm, these linkers are specifically associated with sites of trans-mitochondrial cristae alignment **(Fig. 1k-l)**; in the pig and in the horse, they are positioned at regularly spaced intervals along inter-mitochondrial junctions **(Fig. 1h-j)**. Electron-dense inter-mitochondrial junctions were also seen in cardiomyocytes by classical EM (Duvert et al., 1985; Huang et al., 2013; Picard et al., 2015). Thus, it is plausible that the as-yet-unidentified linker complexes we visualize here represent a more general structural mechanism for orchestrating inter-mitochondrial communication in various cell types.

### Ordered protein arrays at the mitochondria-cytoskeleton interface are conserved across species

To determine how mitochondria interact with the flagellar cytoskeleton, we imaged the mitochondria-cytoskeleton interface in cryo-FIB milled lamellae **(Fig. 2)**. We found that the axoneme-facing surface of the OMM is characterized by an ordered protein array that is absent from the plasma membrane-facing surface **(Fig. 2a)**. These arrays are present in all three species and along the entire midpiece **(Fig. S3a-f)** and resemble the particle rows seen on the axoneme-facing surface of the OMM in guinea pig sperm (Friend and Heuser, 1981) and in mouse sperm (Woolley et al., 2005) by freeze-fracture EM. We observed direct interactions between the arrays and cytoskeletal filaments surrounding the ODFs **(Fig. 2b-c)**, indicating that these arrays tether mitochondria to the midpiece cytoskeleton.

**Fig. 2.**
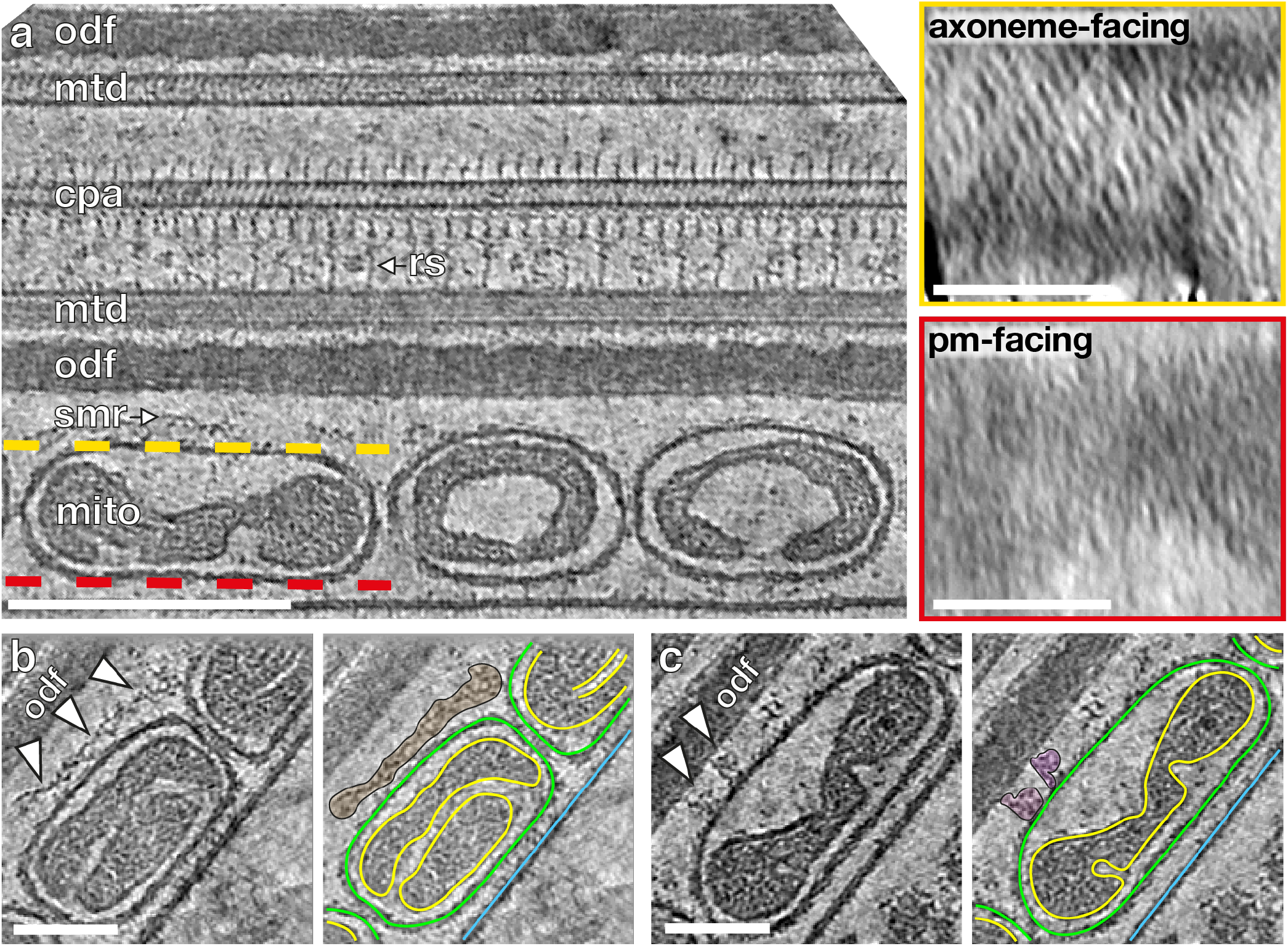
Ordered protein arrays on the outer mitochondrial membrane directly interact with the submitochondrial reticulum. **(a)** Slice through a cryo-tomogram of a FIB-milled horse sperm midpiece, showing mitochondria (mito), the submitochondrial reticulum (smr) outer dense fibers (odf), microtubule doublets (mtd), and the central pair apparatus (cpa). Note how individual complexes (like the radial spoke, rs) are visible in the raw tomogram. The ordered protein array is only found on the axoneme-facing surface (yellow) of midpiece mitochondria, and not on the plasma membrane-facing surface (red). **(b,c)** Slices through a cryo-tomogram of a FIB-milled horse sperm midpiece showing how the array directly interacts with the submitochondrial reticulum to anchor mitochondria to the ciliary cytoskeleton (arrowheads). In right panels, the outer mitochondrial membrane is traced in green, the inner mitochondrial membrane in yellow, and the plasma membrane in blue. **Scale bars:** (a) left – 250 nm, insets – 100 nm; (b,c) 100 nm.

We then aligned and averaged sub-volumes containing the protein arrays and the underlying OMM **(Fig. 3, Table S1)**. Our averages revealed ~22-nm-long two-fold-symmetric boat-shaped structures connected via four densities to a porous membrane **(Fig. 3, Fig. S3g-i)**. Each boat-shaped particle rises ~5 nm above the membrane and consists of two tilde-shaped densities arranged end-to-end. The boat-shaped structures form rows in which each particle is related to its closest neighbors by a ~10 nm translation perpendicular to the particle long axis and a ~6 nm shift along this axis, yielding a center-to-center spacing of ~12 nm **(Fig. 3d-f)**. Each row is oriented ~120° to the long axis of the flagellum and adjacent rows are spaced ~12 nm apart, forming extensive arrays on the axoneme-facing surface of the OMM **(Fig. 3g)**. Remarkably, the averages we obtained from the three species were highly similar, both in terms of individual particle dimensions and in terms of their supramolecular arrangement **(Fig. 3, Fig. S3)**. This conservation suggests that these arrays are a crucial structural element of the mitochondrial sheath.

**Fig. 3.**
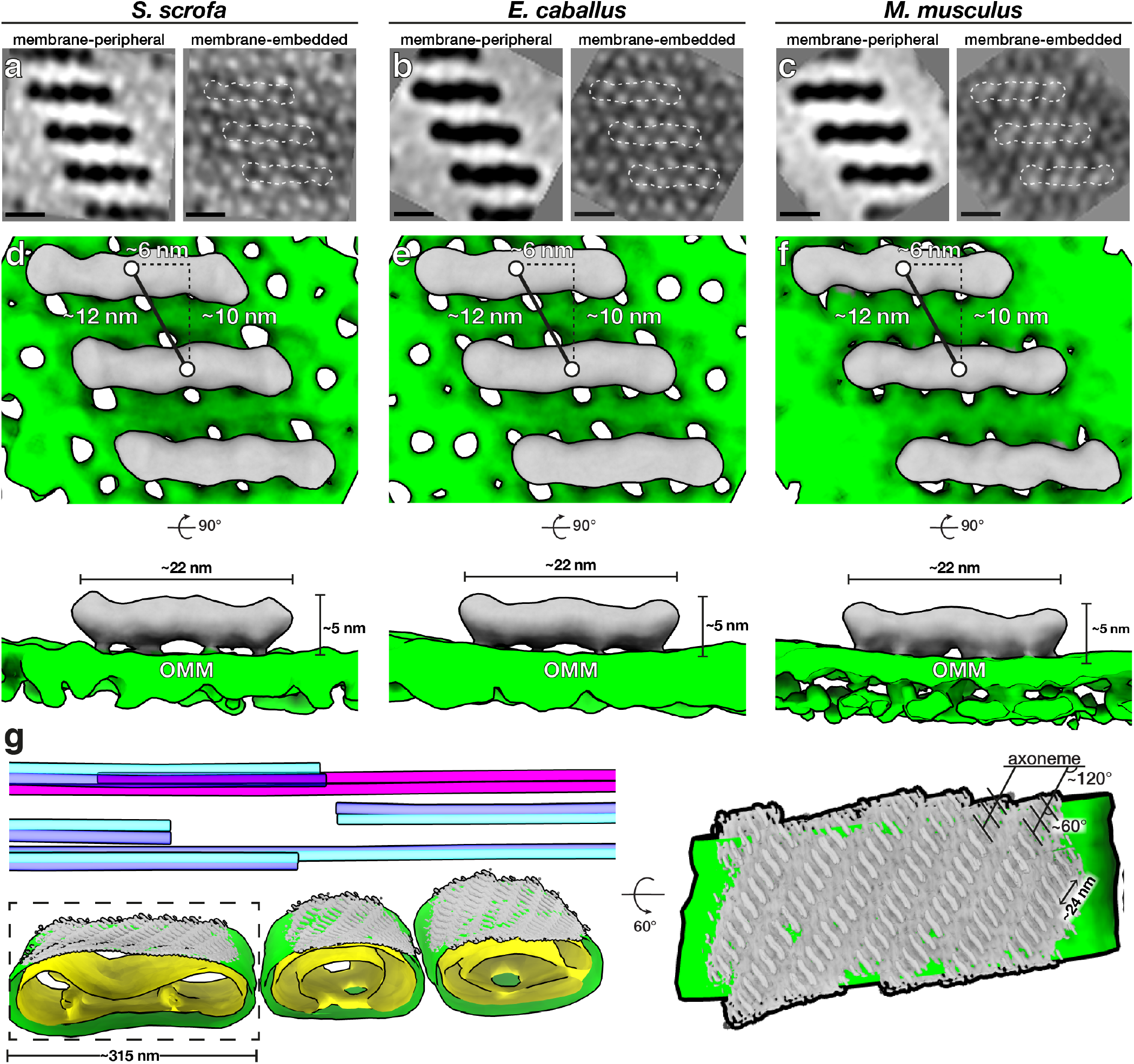
Ordered protein arrays at the mitochondria-cytoskeleton interface are conserved across species. **(a-c)** Subtomogram averages of the protein arrays and underlying outer mitochondrial membrane (OMM) after applying twofold symmetry (note that density is black). **(d-f)** Isosurface renderings of the subtomogram averages in (a-c) with boat-shaped particles in grey and the OMM in green. **(g)** Left panel: Segmentation of the tomogram shown in Figure 2a, with the OMM in green, the IMM in yellow, microtubule doublets in blue, and the cpa in pink. Subtomogram averages of boat-shaped particles are colored grey and plotted back into their positions and orientations in the tomogram. Right panel: Rotated and zoomed-in view of the axoneme-facing surface of a mitochondrion. The axoneme is oriented horizontally, so the ladder-like arrays are oriented ~120°to the flagellar long axis, and individual particles within the array are oriented ~60°to this axis. **Scale bars:** (a-c) 10 nm.

Our averages revealed that the OMM underlying the protein arrays is studded with ~3-4 nm pores arranged in a pseudo-lattice with a center-to-center spacing of ~5 nm. **(Fig. 3a-c, Fig. S3g-i)**. These pore sizes are consistent with the diameters of the voltage dependent anion channels (VDACs), which are known to form ordered arrays in the OMM (Gonçalves et al., 2007; Guo and Mannella, 1993; Hoogenboom et al., 2007; Mannella, 1982). Indeed, our label-free quantitative proteomics experiments show that the most abundant OMM proteins in pig sperm are VDAC2 and VDAC3 **(Table S2)**. Furthermore, the lattice dimensions in our averages closely match those of VDAC in purified Neurospora OMM (Guo and Mannella, 1993; Mannella, 1998). The lattice can be modeled by fitting multiple copies of the VDAC2 crystal structure (Schredelseker et al., 2014) **(Fig. S4a)**. We oriented VDAC2 in the membrane plane based on its known topology (Bayrhuber et al., 2008; Tomasello et al., 2013); however, at the current resolution, we cannot determine the orientation around the pore axis. Thus, in our model, each boat-shaped particle stretches across 8 VDAC molecules **(Fig. 3)**.

### Glycerol kinase-like proteins are probable constituents of the conserved arrays at the mitochondria– cytoskeleton interface

To search for possible constituents of the protein arrays on the VDAC lattice, we used in-cell cross-linking mass spectrometry (XL-MS) (Fasci et al., 2018; Liu et al., 2018) to find potential VDAC2/VDAC3 interaction partners on the OMM **(Fig. 4)**. We treated pig sperm cells with the cross-linker disuccinimidyl sulfoxide (DSSO), which covalently links free lysines that are within ~3 nm (Cα-Cα) of each other. To increase confidence, we screened for cross-links identified with at least two cross-link spectral matches (CSMs) (see Materials and Methods for details).

**Fig. 4.**
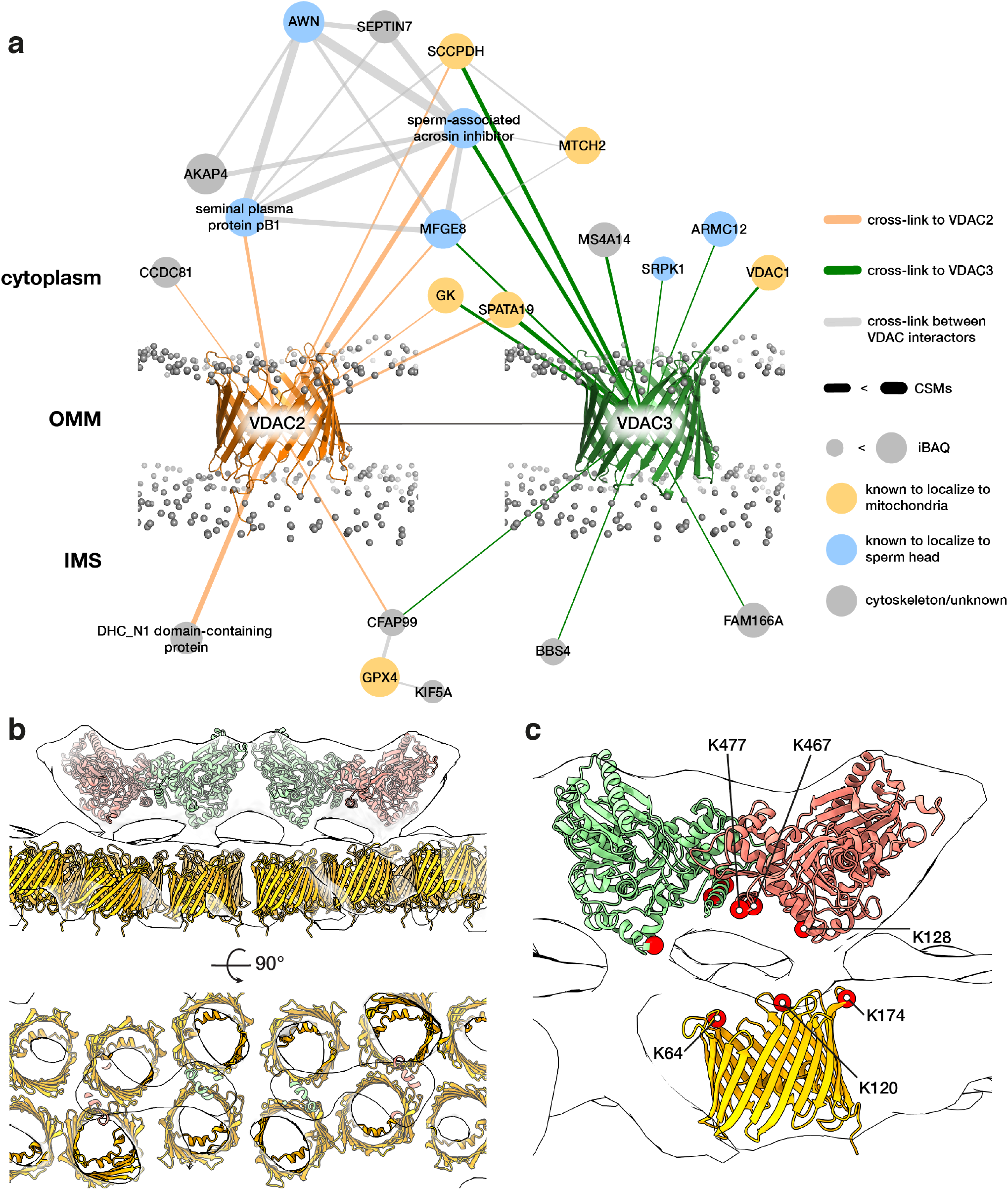
Modelling the outer mitochondrial membrane (OMM) array as glycerol kinase-like (GK) proteins anchored on voltage dependent anion channels (VDACs). **(a)** The VDAC2/VDAC3 interactome derived from in-cell XL-MS of pig sperm. Protein nodes are colored according to their known subcellular localizations (yellow – mitochondria, blue – head, grey – cytoskeleton/unknown). Gray spheres indicate the phosphate groups of a simulated lipid bilayer which was structurally aligned based on the simulation for monomeric mouse VDAC1 (PDB 4C69) obtained from the MemProtMD server (Newport et al., 2019). **(b)** Modeling the OMM array as GK-like proteins anchored on VDACs. A GK-like dimer-of-dimers homology model (red and green) and VDAC homology models (yellow) were fitted into the pig subtomogram average map (white). **(c)** The positions of cross-linked Lys residues (red circles) are consistent with GK and VDAC orientation assignments in our model.

We first screened candidate proteins based on their known subcellular localizations **(Fig. 4a)**. VDAC2/VDAC3 cross-linked to mitochondria-associated proteins as well as to sperm head-associated proteins. This is consistent with immunofluorescence studies localizing VDAC2/VDAC3 both to the midpiece and to the acrosome, a large vesicle capping the anterior sperm nucleus (Hinsch et al., 2004; Kwon et al., 2013). Of the proteins in the mitochondria-associated interaction hub, three proteins are particularly noteworthy because they are known to localize to the OMM and because their disruption results in dysplasia of the mitochondrial sheath: spermatogenesis-associated protein 19 (SPATA19) (Mi et al., 2015), glutathione peroxidase 4 (GPX4) (Imai et al., 2009; Schneider et al., 2009), and glycerol kinase (GK) (Chen et al., 2017b; Shimada et al., 2019).

To distinguish among these candidates, we compared the location of the cross-links with the known topology of VDAC in the OMM (Bayrhuber et al., 2008; Tomasello et al., 2013). GPX4 would interact on the side facing the intermembrane space, whereas SPATA19 and GK would interact on the cytoplasmic face. Both SPATA19 and GK are highly abundant **(Table S2)**, as would be expected for proteins forming extensive arrays. Assuming an average protein density of ~1.43 g/cm^3^ (Quillin and Matthews, 2000), which corresponds to ~0.861 Da/Å^3^, we estimate that each boat-shaped particle in the array has a molecular weight of ~250 kDa. SPATA19 is a small protein with an estimated molecular weight of ~18 kDa. To fit into our EM densities, it must either be present in multiple copies or form a complex with other proteins. In contrast, GK has an estimated molecular weight of ~60 kDa and is known to form S-shaped dimers (~120 kDa) that are conserved from bacteria (Bystrom et al., 1999; Fukuda et al., 2016) to eukaryotes (Balogun et al., 2019; Schnick et al., 2009).

To build a GK-VDAC model based on our subtomogram average, we used rigid-body fitting to place two GK dimers end-to-end into a boat-shaped density **(Fig. S4b, Fig. 4b)**. These fits defined a clear orientation for GK, with the Ntermini pointing upwards and the C-terminal helices facing the OMM **(Fig. 4b)**. To validate our fits, we mapped the cross-linked lysines onto the resulting model **(Fig. 4c)**. All cross-links were between the cytosolic face of VDAC2 and the OMM-facing surface of GK, which is consistent with the orientation expected from our fits. Assigning GK-like proteins as constituents of the ordered OMM arrays at the mitochondria-cytoskeleton interface is also supported by recent genetic studies. Sperm from mice lacking sperm-specific GK isoforms, which do not show glycerol kinase activity *in vitro* (Pan et al., 1999), have disorganized mitochondrial sheaths (Chen et al., 2017b; Shimada et al., 2019). In these mice, spherical mitochondria properly align along the flagellum but fail to properly elongate and coil around the ODFs (Shimada et al., 2019). This phenotype is consistent with our data showing direct links between GK protein arrays and the submitochondrial reticulum **(Fig. 2b-c)**.

## Discussion

In this study, we used cryo-FIB milling-enabled cryoET to image the sperm mitochondrial sheath in three mammalian species. Our data reveal that overall mitochondrial dimensions are remarkably consistent in sperm from the same species **(Fig. 1, S1)**. This contrasts with other mitochondria-rich tissues such as skeletal muscle, where there are massive variations in mitochondrial size and morphology within individual cells (Vincent et al., 2019). In addition, we did not observe mitochondrial nanotunnels in any of the species we examined, in contrast to their relative abundance in muscle tissue (Vincent et al., 2017, 2019). Our data also show that mitochondrial dimensions and cristae architecture vary across species **(Fig. 1)**, providing possible structural bases for interspecific differences in mitochondrial energetics. Further comparative studies of how mitochondrial structure varies with sperm metabolism will undoubtedly contribute to our broader understanding of how mitochondrial form relates to function.

Our data show that, despite this diversity, the molecular underpinnings of mitochondrial sheath architecture are conserved, at least in mammals. Specifically, we identified novel inter-mitochondrial linkers that tether adjacent mitochondria **(Fig. 1, S2)** and arrays of boat-shaped particles that anchor mitochondria to the cytoskeleton **(Fig. 2, 3)**. In-cell subtomogram averaging and in-cell XL-MS suggest that these arrays consist of GK-like proteins anchored on VDAC lattices in the OMM **(Fig. 4)**. Given that VDACs are ubiquitous OMM proteins, our findings motivate further efforts to explore whether they also regulate mitochondria-cytoskeleton interactions in other cell types.

The OMM arrays may function to regulate the precise elongation and coiling of mitochondria, contributing to the striking consistency within the mitochondrial sheath. In mature sperm, these arrays may help maintain the integrity of mitochondria-cytoskeleton contacts, stabilizing them against shear stresses during sperm motility and hyperactivation. However, it is unclear what determines the organization of these arrays in the first place. Our averages do not hint at direct interactions between boat-shaped particles. Instead, their spacing may be defined by the organization of the underlying VDAC lattice. Another intriguing possibility is that the arrays are organized by their cytoskeletal binding partners; the periodicity of relevant motifs on the submitochondrial reticulum could dictate the spacing of the OMM arrays.

To our knowledge, this is the first time such assemblies have been visualized at any organelle-cytoskeleton interface in any cell type. Defining whether similar arrays are present in other differentiated cell types – and whether they use a similar pool of protein components – is an area ripe for study. In striated muscle, proper mitochondrial positioning is critical for muscle function and depends on direct associations between mitochondria and intermediate filaments (Konieczny et al., 2008; Milner et al., 2000). Similarly, in skin cells, mitochondrial organization depends on keratin (Steen et al., 2020). The structural bases for these associations are unknown, but cryo-ET and in-cell XL-MS may prove useful in these contexts as well.

## Acknowledgements

The authors thank Dr. M Vanevic for excellent computational support; Dr. D Vasishtan for providing scripts that greatly facilitated subtomogram averaging; Ingr. CTWM Schneijdenberg and JD Meeldijk for managing and maintaining the Utrecht University EM Square facility; Stal Schep (Tull en het Waal, The Netherlands) for providing horse semen; MW Haaker and M Houweling for providing mouse reproductive tracts; Prof. F Förster and Prof. A Akhmanova for critical reading of the manuscript; and Prof. EY Jones for insightful discussions. The authors also thank the Henriques Lab for the publicly-available LATEX template. This work benefitted from access to the Netherlands Center for Electron Nanoscopy (NeCEN) with support from operators Dr. RS Dillard and Dr. CA Diebolder and IT support from B Alewijnse. RZC, JFH, and AJRH acknowledge support from NWO funding the Netherlands Proteomics Centre through the X-omics Road Map program (project 184.034.019). This work was funded by NWO Start-Up Grant 740.018.007 to TZ, and MRL is supported by a Clarendon Fund-Nuffield Department of Medicine Prize Studentship.

## Author Contributions

PM, MZ, HH, EGB, and BMG provided sperm samples. MRL, MCR, and RTR prepared samples for cryo-ET. MRL performed cryo-FIB milling. MRL, MCR, RTR, SCH, and TZ collected cryo-ET data. MRL and MCR processed cryo-ET data. MRL, MCR, and TZ analyzed cryo-ET data. RZC and JFH performed all proteomics and cross-linking mass spectrometry experiments along with corresponding structural modelling under the supervision of AJRH. MRL and TZ wrote the manuscript, and all authors contributed to revisions.

## Declaration of Interests

The authors declare no competing interests.

## Materials and Methods

### Sperm collection and preparation

Pig sperm samples were purchased from an artificial insemination company (Varkens KI Nederland), stored at 18°C, and prepared for imaging within 1 day of delivery. Sperm were layered onto a discontinuous gradient consisting of 4 mL of 35% Percoll^®^ (GE Healthcare) underlaid with 2 mL of 70% Percoll^®^, both in HEPES-buffered saline (HBS: 20 mM HEPES, 137 mM NaCl, 10 mM glucose, 2.5 mM KCl, 0.1% kanamycin, pH 7.6) and centrifuged at 750*g* for 15 min at RT (Harrison et al., 1993). Pelleted cells were washed once in phosphatebuffered saline (PBS: 137 mM NaCl, 3 mM KCl, 8 mM Na_2_HPO_4_, 1.5 mM KH_2_PO_4_, pH 7.4), resuspended in PBS and counted with a hemocytometer.

Horse semen was collected from mature Warmblood stallions using a Hanover artificial vagina in the presence of a teaser mare. After collection, semen was filtered through gauze to remove gel fraction and large debris, then transported to the laboratory at 37°C and kept at room temperature until further processing. Semen was diluted in INRA96 (IMV Technologies) to obtain a sperm concentration of 30 × 10^6^ cells/mL. After this, sperm were centrifuged through a discontinuous Percoll gradient as described above for pig sperm for 10 min at 300*g* followed by 10 min at 750*g* (Harrison et al., 1993). The remaining pellet was resuspended in 1 mL of PBS and centrifuged again for 5 min at 750*g*.

Mouse sperm were collected from the cauda epididymis of adult male C75BL/6 mice as described in (Hutcheon et al., 2017). Briefly, male mice were culled as described in (Mederacke et al., 2015) and the cauda epididymides were dissected with the vas deferens attached and placed in a 500 μL droplet of modified Biggers, Whitten, and Whittingham media (BWW: 20 mM HEPES, 91.5 mM NaCl, 4.6 mM KCl, 1.7 mM D-glucose, 0.27 mM sodium pyruvate, 44 mM sodium lactate, 5 U/mL penicillin, and 5 μg/mL streptomycin, adjusted to pH 7.4 and an osmolarity of 300 mOsm/kg). To retrieve the mature cauda spermatozoa from the epididymides, forceps were used to first gently push the stored sperm from the vas deferens, after which two incisions were made with a razor blade in the cauda. Spermatozoa were allowed to swim out of the cauda into the BWW over a period of 15 min at 37°C, after which the tissue was removed and sperm were loaded onto a 27% Percoll density gradient and washed by centrifugation at 400*g* for 15 min. The pellet consisting of an enriched sperm population was resuspended in BWW and again centrifuged at 400*g* for 2 min to remove excess Percoll.

### Cryo-EM grid preparation

Typically, 3 μL of a suspension containing either 2-3 × 10^6^ cells/mL (for whole cell tomography) or 20-30 × 10^6^ cells/mL (for cryo-FIB milling) was pipetted onto a glow-discharged Quantifoil R 2/1 200-mesh holey carbon grid. One μL of a suspension of BSA-conjugated gold beads (Aurion) was added, and the grids then blotted manually from the back (opposite the side of cell deposition) for ~3 s (for whole cell tomography) or for ~5-6 s (for cryo-FIB milling) using a manual plunge-freezer (MPI Martinsreid). Grids were immediately plunged into a liquid ethane-propane mix (37% ethane) (Tivol et al., 2008) cooled to liquid nitrogen temperature. Grids were stored under liquid nitrogen until imaging.

### Cryo-focused ion beam milling

Grids were mounted into modified Autogrids (ThermoFisher) for mechanical support. Clipped grids were loaded into an Aquilos (ThermoFisher) dual-beam cryo-focused ion beam/scanning electron microscope (cryo-FIB/SEM). All SEM imaging was performed at 2 kV and 13 pA, whereas FIB imaging for targeting was performed at 30 kV and 10 pA. Milling was typically performed with a stage tilt of 18°, so lamellae were inclined 11° relative to the grid. Each lamella was milled in four stages: an initial rough mill at 1 nA beam current, an intermediate mill at 300 pA, a fine mill at 100 pA, and a polishing step at 30 pA. Lamellae were milled with the wedge pre-milling technique described in (Schaffer et al., 2017) and with expansion segments as described in (Wolff et al., 2019).

### Tilt series acquisition

Tilt series were acquired on either a Talos Arctica (ThermoFisher) operating at 200 kV or a Titan Krios (ThermoFisher) operating at 300 kV, both equipped with a post-column energy filter (Gatan) in zero-loss imaging mode with a 20-eV energy-selecting slit. All images were recorded on a K2 Summit direct electron detector (Gatan) in either counting or super-resolution mode with dose-fractionation. Tilt series were collected using SerialEM (Mastronarde, 2005) at a target defocus of between −4 and −6 μm (conventional defocus-contrast) or between −0.5 and −1.5 μm (for tilt series acquired with the Volta phase plate). Tilt series were typically recorded using either strict or grouped dose-symmetric schemes, either spanning ± 56° in 2° increments or ± 54° in 3° increments, with total dose limited to ~100 *e^-^*/Å^2^.

### Tomogram reconstruction

Frames were aligned either post-acquisition using Motioncor2 1.2.1 (Zheng et al., 2017) or on-the-fly using Warp (Tegunov and Cramer, 2019). Frames were usually collected in counting mode; when super-resolution frames were used, they were binned 2X during motion correction. Tomograms were reconstructed in IMOD (Kremer et al., 1996) using weighted back-projection, with a SIRT-like filter (Zeng, 2012) applied for visualization and segmentation. Defocus-contrast tomograms were CTF-corrected in IMOD using *ctfphaseflip* while VPP tomograms were left uncorrected.

### Tomogram segmentation

Segmentation was generally performed semi-automatically using the neural network-based workflow implemented in the TomoSeg package in EMAN 2.21 (Chen et al., 2017). Microtubules, however, were traced manually in IMOD. Segmentation was then manually refined in Chimera 1.12 (Pettersen et al., 2004) or in ChimeraX (Goddard et al., 2018). Visualization was performed in ChimeraX.

### Subtomogram averaging of ATP synthase and outer mitochondrial membrane arrays

Subtomogram averaging with missing wedge compensation was performed using PEET 1.13.0 (Heumann et al., 2011; Nicastro et al., 2006). Resolution was estimated using the Fourier shell correlation (FSC) at a cut-off of 0.5 (Nicastro et al., 2006). Alignments were generally performed first on binned data, after which aligned positions and orientations were transferred to less-binned data using scripts generously provided by Dr. Daven Vasishtan. Details of acquisition parameters and particle numbers are summarized in.. Table S1.

Alignment strategies for these complexes were designed to take advantage of their defined orientations relative to the membrane plane. Particles were picked manually and their initial orientations were defined using *stalkInit*. Initial references were either a randomly chosen particle (for ladder-like arrays) or an average of all particles after roughly aligning them based on their initial orientations (for ATP synthase). Independent alignments using independent initial references were performed for datasets from different species. Alignments allowed for large rotational search ranges around the particle long axis (defined as the y-axis, perpendicular to the membrane plane), with limited search ranges around the x- and z-axes (the membrane plane).

All initial alignments were performed without symmetry. After visual inspection of the maps, twofold symmetry was applied for ladder-like arrays. Symmetrization involved using the aligned positions from the unsymmetrized runs as seed points and rotating particles around the axis of symmetry to generate virtual particles. A symmetrized volume was generated by averaging all particles and virtual particles and used as a reference for a final, restricted alignment.

Plotbacks were generated in IMOD by first running *createAlignedModel* to generate model files reflecting updated particle positions and orientations after alignment. The relevant subtomogram average was then thresholded for visualization and saved as an isosurface model, which was then placed back into the tomograms using *clonemodel*.

### Measurements and quantification

All measurements of mitochondrial width were performed in IMOD on Volta phase plate tomograms filtered with a SIRT-like filter. Mitochondrial width was measured in the non-missing wedge direction at five points along the length of each mitochondrion. Only mitochondria that were entirely in the field of view were included in the measurements. Tomograms and corresponding measurements were then grouped based on their locations relative to the connecting piece, which were determined based on low-magnification images used for targeting during data acquisition.

Internal mitochondrial ultrastructure was quantified from tomograms from cryo-FIB milled lamellae. The volume occupied by the matrix (V_matrix_, the volume enclosed by the IMM) was measured relative to the volume occupied by the entire mitochondrion (V_mito_, the volume enclosed by the OMM). Mesh volumes were extracted from segmentations using *imodinfo*. Because neural network-based segmentation often resulted in gaps, mitochondrial membranes were segmented manually in IMOD for quantification. Only slices in which both the IMM and OMM were clearly defined were used for segmentation.

### Cross-linking, lysis, digestion and peptide fractionation

All proteomics and cross-linking mass spectrometry ex-periments were performed on Percoll-washed pig sperm prepared as described above. For cross-linking, approximately 300 × 106 cells were used from 3 different animals. Briefly, pelleted sperm cells were resuspended in 540 μL of PBS supplemented with disuccinimidyl sulfoxide (DSSO, Thermo Fisher Scientific) to a final concentration of 1 mM. The reaction mix was incubated for 30 min at 25°C with 700 rpm shaking in a ThemoMixer C (Eppendorf) and subsequently quenched for 20 min by adding Tris-HCl (final concentration 50 mM). Cross-linked cells were spun down at 13 800g for 10 min at 4°C, after which the supernatant was removed. Cells were then lysed according to a protocol modified from (Potel et al., 2018). Cells were resuspended in 1 mL of lysis buffer (100 mM Tris-HCl pH 8.5, 7 M Urea, 1% Triton X-100, 5 mM TCEP, 30 mM CAA, 10 U/ml DNase I, 1 mM MgCl2, 1% benzonase (Merck Millipore, Darmstadt, Germany), 1 mM sodium orthovanadate, phosphoSTOP phosphatases inhibitors, and cOmpleteTM Mini EDTA-free protease inhibitors). Cells were sonicated on ice for 2 min using an ultrasonic processor (UP100H, Hielscher) at 80% amplitude. The proteins were then precipitated according to (Wessel and Flügge, 1984) and the dried protein pellet resuspended in digestion buffer (100 mM Tris-HCl pH 8.5, 1% sodium deoxycholate (Sigma-Aldrich), 5 mM TCEP, and 30 mM CAA). Trypsin and Lys-C proteases were added to a 1:25 and 1:100 ratio (w/w) respectively and protein digestion performed overnight at 37°C. The final peptide mixtures were desalted with solid-phase extraction C18 columns (Sep-Pak, Waters) and fractionated with an Agilent 1200 HPLC pump system (Agilent) coupled to a strong cation exchange (SCX) separation column (Luna SCX 5 μm – 100 Å particles, 50 × 2mm, Phenomenex), resulting in 25 fractions.

### Liquid chromatography with mass spectrometry

Ap-proximately 1000 ng of peptides from each biological replicate before SCX fractionation were first injected onto an Agilent 1290 Infinity UHPLC system (Agilent) on a 50-cm analytical column packed with C18 beads (Dr Maisch Reprosil C18, 3 μm) coupled online to an Orbitrap HF-X (Thermo Fisher Scientific). For this classical bottom-up analysis, we used the following LC-MS/MS parameters: after 5 min of loading with 100% buffer A (H2O with 0.1% formic acid), peptides were eluted at 300 nL/min with a 95-min gradient from 13% to 40% of buffer B (80% acetonitrile and 20% H2O with 0.1% formic acid). For MS acquisition we used an MS1 Orbitrap scan at 60,000 resolution, automatic gain control (AGC) target of 3 × 106 ions and maximum inject time of 20 ms from 375 to 1600 m/z; the 15 most intense ions were submitted to MS2 Orbitrap scan at 30,000 resolution, AGC target of 1 × 105 ions and maximum inject time of 50 ms (isolation window of 1.4 m/z, NCE at 27% and dynamic exclusion of 16 seconds). The SCX fractions were analyzed with same Agilent HPLC and the same nano-column coupled on-line to an Orbitrap Lumos mass spectrometer (ThermoFisher Scientific). For these runs, we used a gradient from 6% to 39% buffer B over 100 min with specific MS settings for DSSO cross-links: survey MS1 Orbitrap scan at 60,000 resolution from 375 to 1500, AGC target of 4 × 105 ions and maximum inject time of 50 ms; MS2 Orbitrap scan at 30,000 resolution, AGC target of 5 × 104 ions, and maximum inject time of 100 ms for detection of DSSO signature peaks (difference in mass of 37.972 Da). The four ions with this specific difference were analyzed with a MS3 Ion Trap scans at AGC target of 2 × 104 ions, maximum inject time of 150 ms for sequencing selected signature peaks (representing the individual peptides).

### Mass spectrometry data processing

The 3 raw files obtained with classical bottom-up approach were analyzed with MaxQuant version 1.6.17 with all the automatic settings adding Deamidation (N) as dynamic modification against the Sus scrofa reference proteome (Uniprot version of 08/2020 with 49,795 entries). With this search, we were able to calculate intensity-based absolute quantification (iBAQ) values and created a smaller FASTA file to use for analysis of cross-linking experiments (table as Supplementary information). Raw files for cross-linked cells were analyzed with Proteome Discoverer software suite version 2.4.1.15 (ThermoFisher Scientific) with the incorporated XlinkX node for analysis of cross-linked peptides as described in (Klykov et al., 2018). Data were searched against the smaller FASTA created in house with “MS2_MS3 acquisition strategy”. For the XlinkX search, we selected full tryptic digestion with 3 maximum missed cleavages, 10 ppm error for MS1, 20 ppm for MS2, and 0.5 Da for MS3 in Ion Trap. For modifications, we used static Carbamidomethyl (C) and dynamic Oxidation (M), Deamidation (N) and Met-loss (protein N-term). The crosslinked peptides were accepted with a minimum score of 40, minimum score difference of 4 and maximum FDR rate set to 5%; further standard settings were used.

### Interactome analysis, homology modelling, and cross-link mapping

The interaction map for VDAC proteins was generated in R (Grant et al., 2006) using the igraph package (v 1.2.4.2). Only cross-links with at least two crosslink spectral matches (CSMs) were used for network generation. Homology models of GK and VDAC2 were generated in Robetta (Kim et al., 2004) and fitted into subtomogram average maps by rigid body fitting in Chimera X. Cross-links were mapped onto the resulting models using ChimeraX.

### Data availability

Subtomogram average maps have been deposited to the Electron Microscopy Data Bank (EMDB) with the following accession numbers: EMDB-12354, 12355, 12356, and 12357. The model of putative glycerol kinase-like proteins anchored on a VDAC array has been deposited to the Protein Data Bank (PDB) with the accession number PDB ID 7NIE.

---

**Fig. S1.**
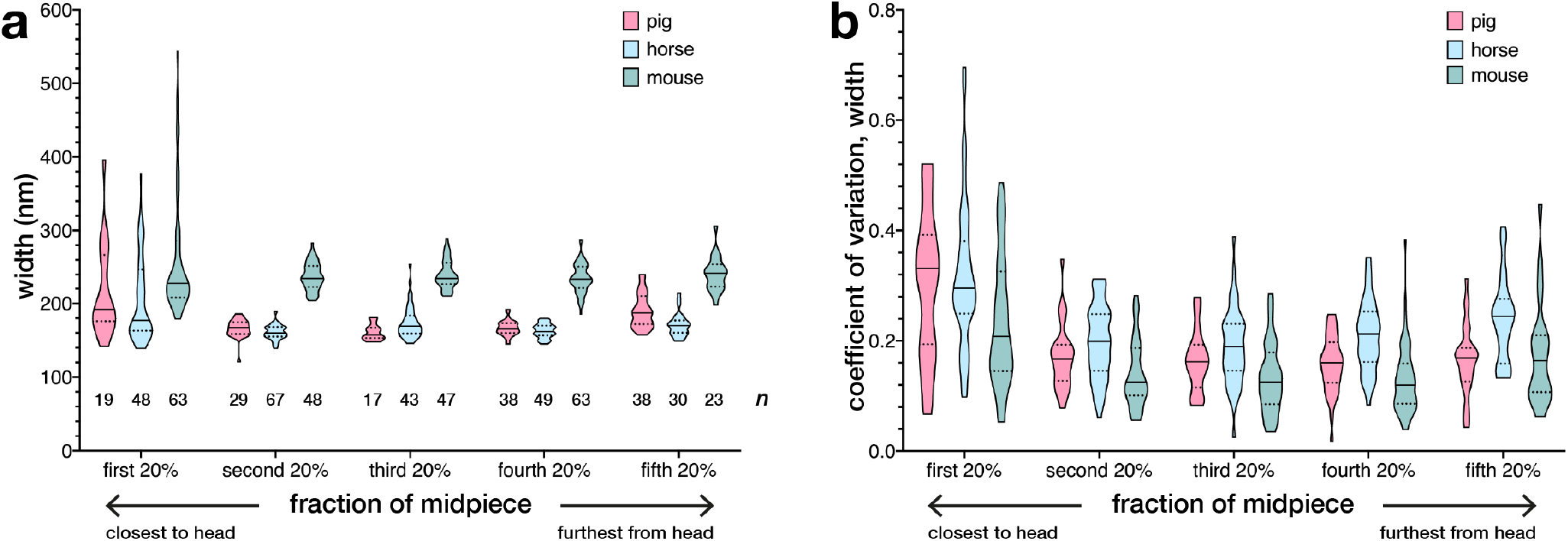
Mitochondrial dimensions are consistent within species but vary across species and spatially along the midpiece. **(a)** Plotting the average width of mitochondria from different regions of the midpiece shows that mouse sperm mitochondria are larger than pig and horse sperm mitochondria. Note also that, in all three species, mitochondria at the proximal end of the midpiece are larger than those in more distal parts. **(b)** Mitochondrial width was measured at five points along the length of each mitochondrion. Plotting the coefficient of variation from different regions along the midpiece shows that mitochondria at the start of the midpiece have more variable widths along their lengths. In (a), n indicates the number of mitochondria analyzed. In both (a) and (b), solid lines represent the median and dotted lines represent the first and third quartiles.

**Fig. S2.**
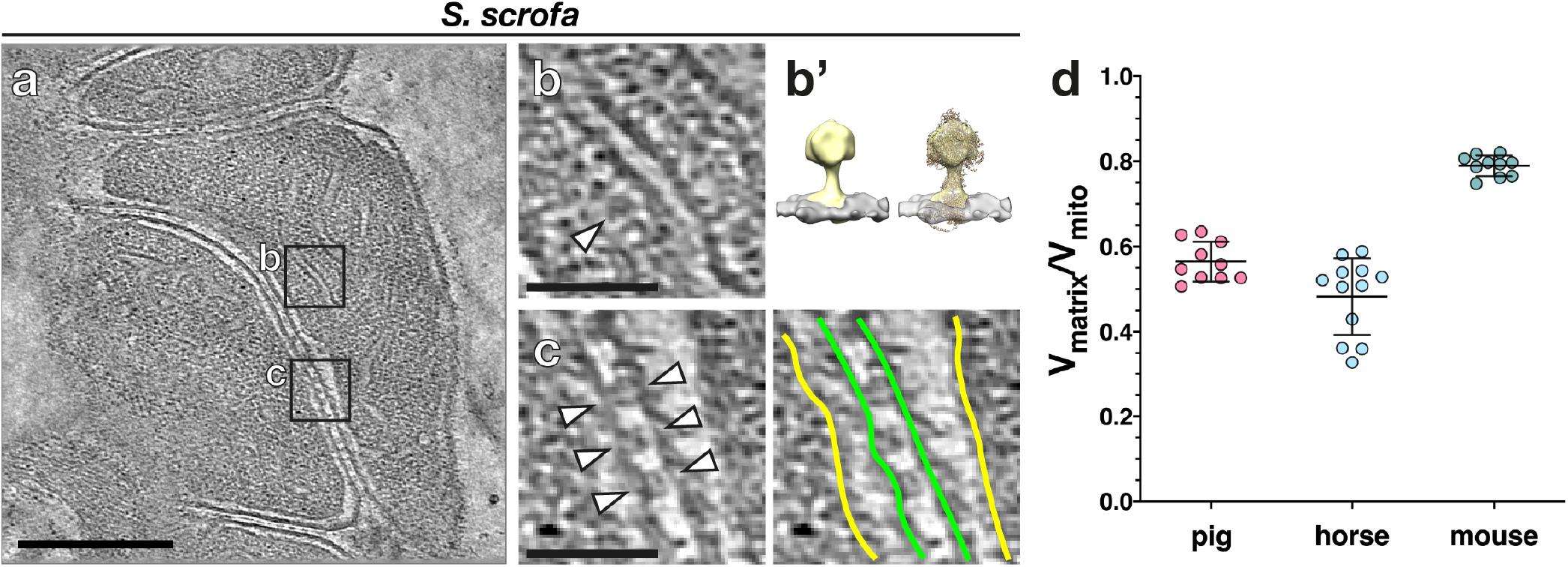
Cryo-focused ion beam (cryo-FIB) milling reveals the internal organization of sperm mitochondria. **(a)** Slice through a cryo-tomogram of FIB-milled pig sperm mitochondria close to the connecting piece. **(b)** ATP synthase can be directly identified on cristae based on its characteristic shape, which is confirmed by subtomogram averaging (b’). **(c)** Novel inter-mitochondrial linkers tether neighboring mitochondria to each other (arrowheads in left panel). **(d)** Quantifying the volume enclosed by the mitochondrial matrix relative to the volume enclosed by the whole mitochondrion reveals that pig and horse sperm mitochondria have more expanded cristae and more condensed matrices than mouse sperm mitochondria. Lines represent mean ± standard deviation.**Scale bars:** (a) 250 nm, (b-c) 100 nm.

**Fig. S3.**
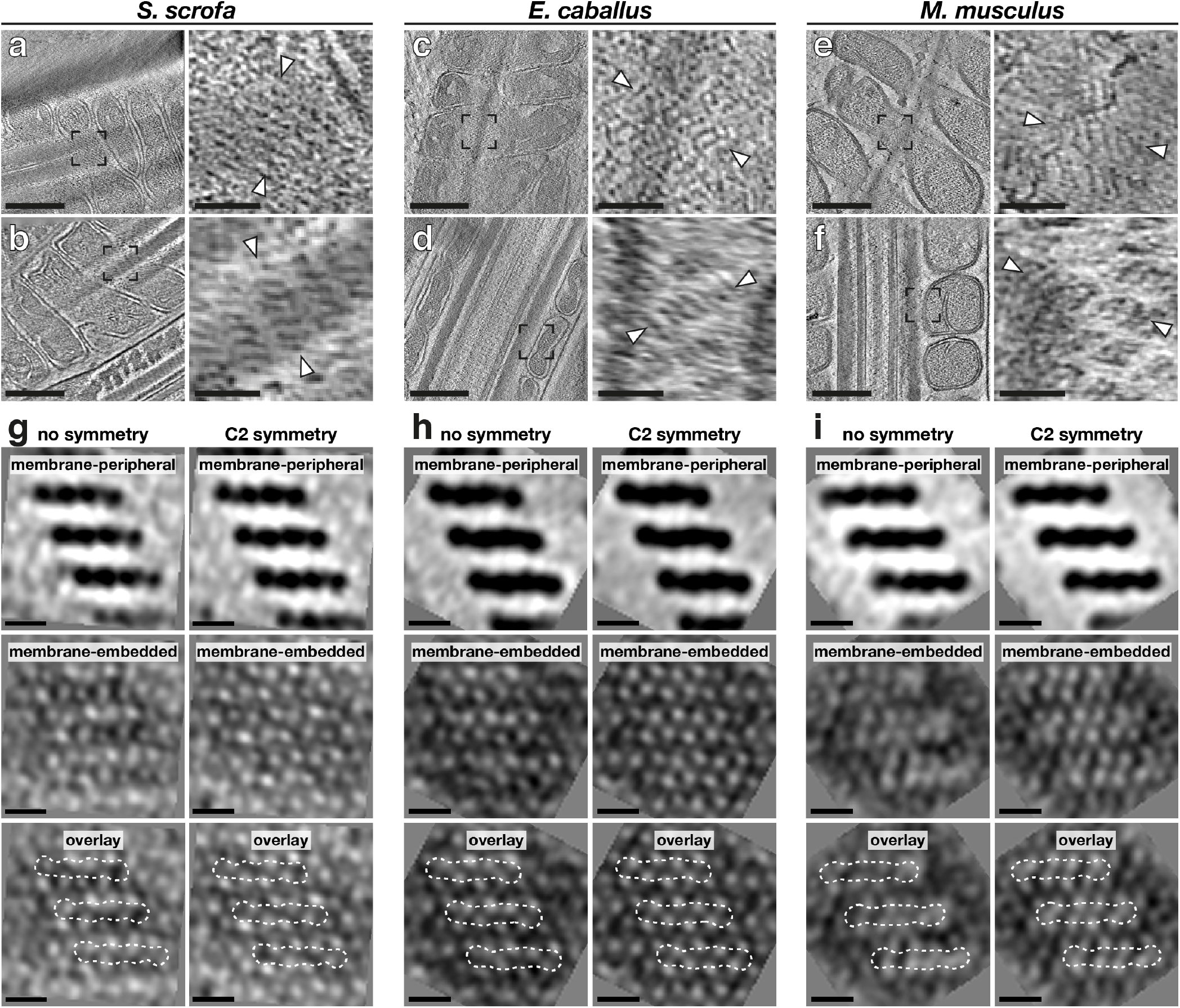
The particles forming the ordered arrays at the mitochondria-cytoskeleton interface are two-fold symmetric. **(a-f)** Slices through cryo-tomograms of FIB-milled pig (a,b), horse (c,d), and mouse (e,f) mitochondria. Right panels show digital zooms of the regions boxed out in the left panels, with arrowheads indicating arrays. **(g-i)** Subtomogram averages of the arrays and the outer mitochondrial membrane (OMM) without (left) and with (right) twofold symmetry. **Scale bars:** (a-f) 250 nm, (g-i) 10 nm.

**Fig. S4.**
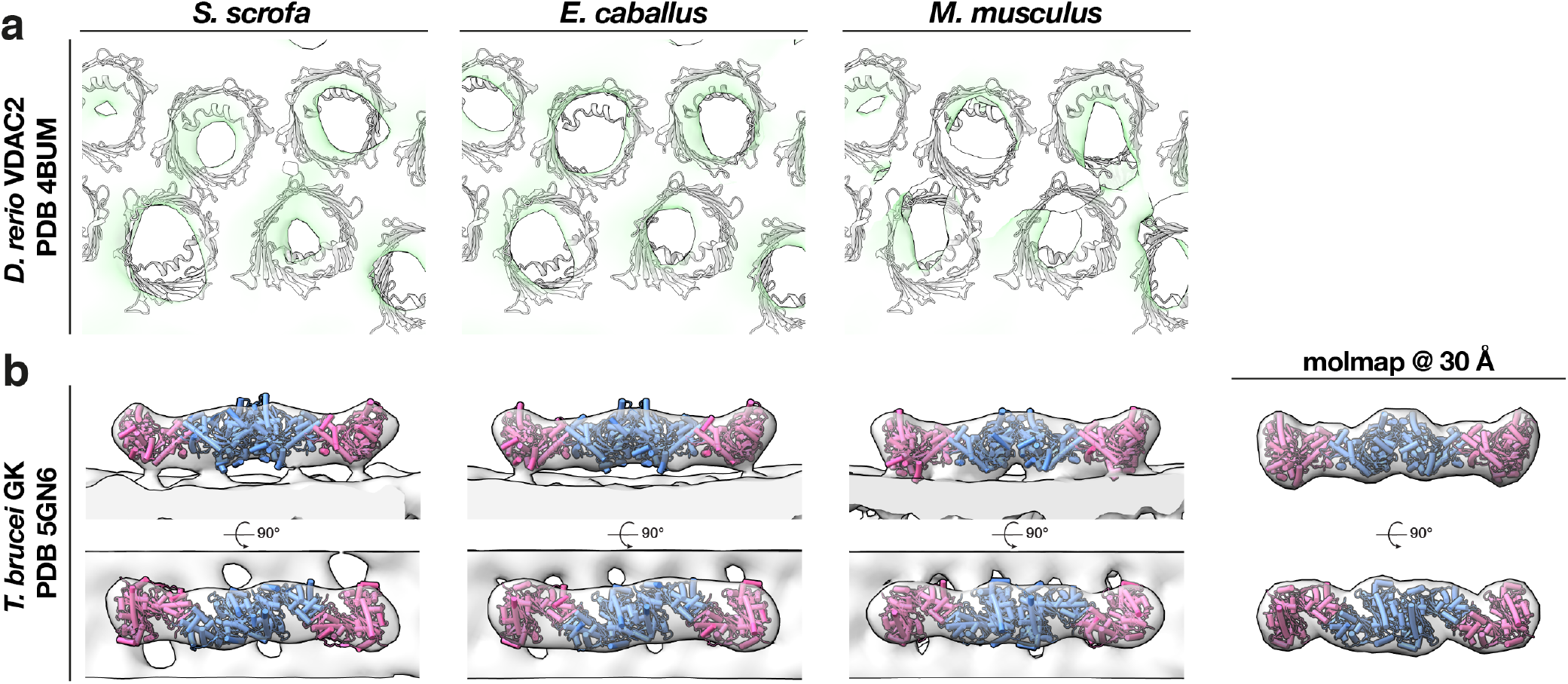
Fitting crystal structures of glycerol kinase (GK and voltage dependent anion channels (VDACs) into the pig subtomogram average map. **(a)** The crystal structure of VDAC2 from zebrafish (PDB 4BUM) is shown in grey, fitted into the cryo-ET averaged map (green). **(b)** Two copies of a crystal structure of GK (pink and blue) from Trypanosoma brucei (PDB 5GN6) fitted into the cryo-ET averaged map (grey). On the right, the GK crystal structure is shown filtered to 30Å resolution.

**Table S1.**
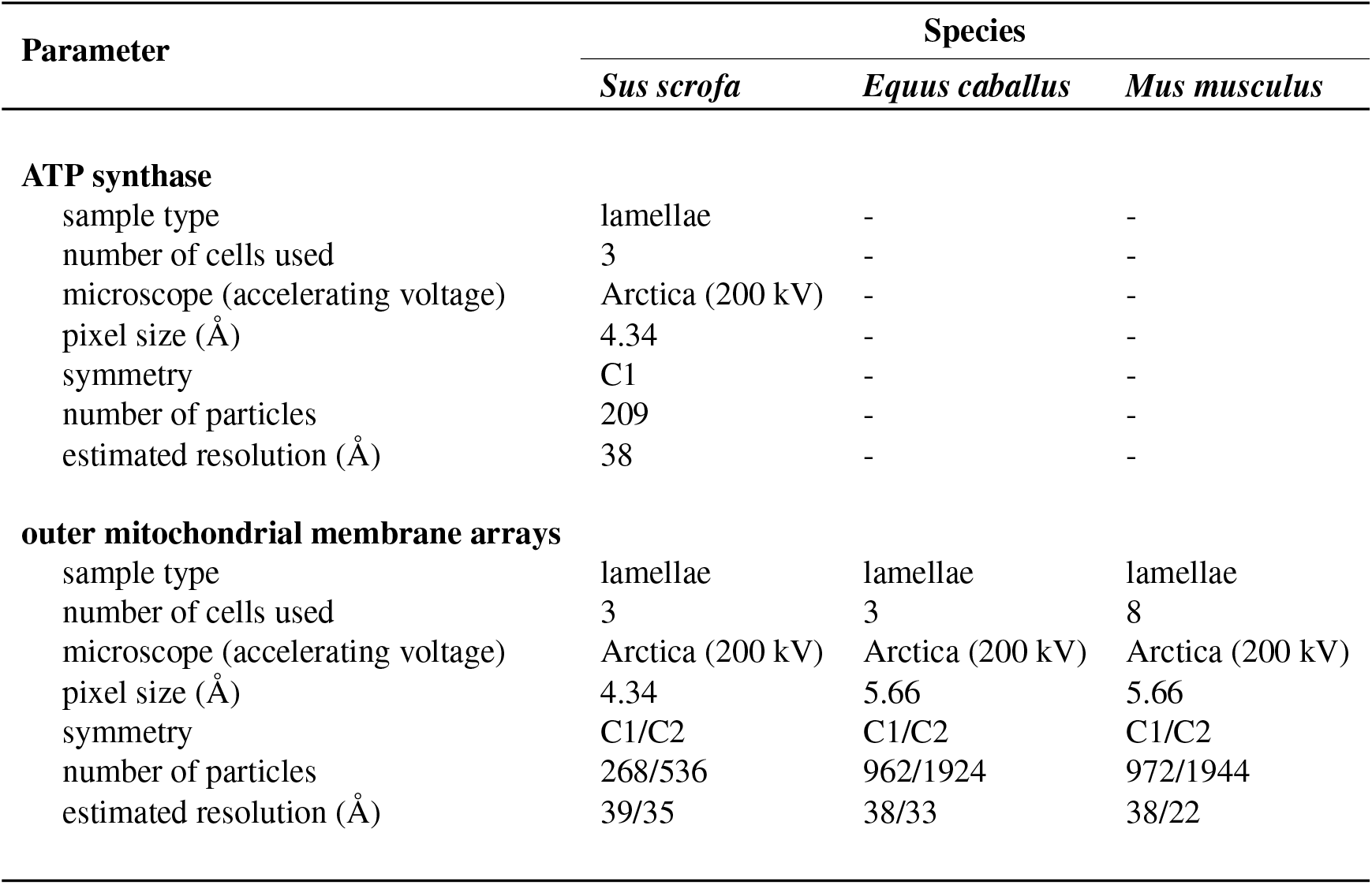
Image acquisition and processing metrics for subtomogram averaging of mitochondrial protein complexes in mammalian sperm.

**Table S2.**
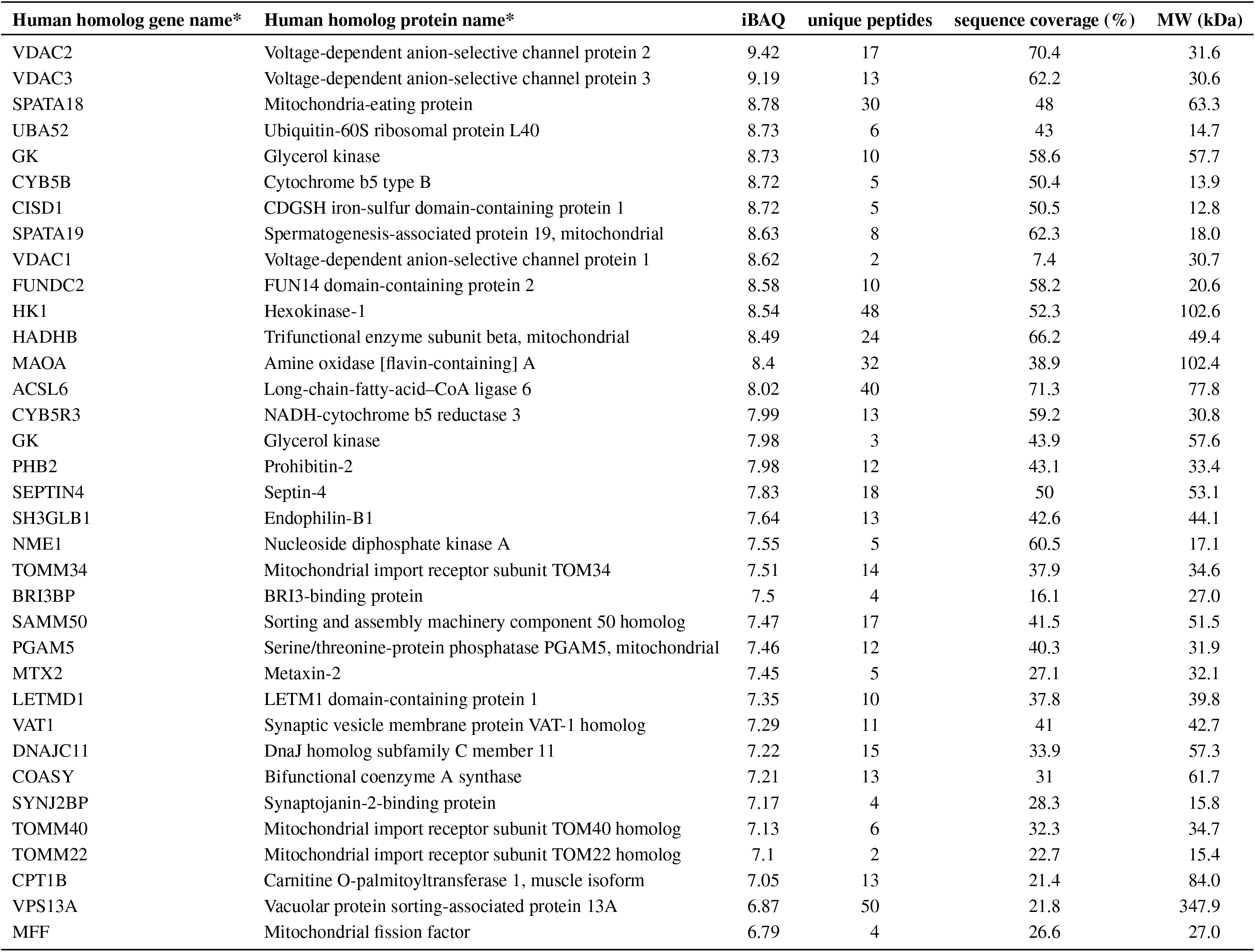
Top 35 most abundant outer mitochondrial membrane proteins identified in the pig sperm proteome.

